# Orsay virus infection increases *Caenorhabditis elegans* resistance to heat-shock

**DOI:** 10.1101/2023.11.02.565305

**Authors:** Victoria G. Castiglioni, Santiago F. Elena

**Affiliations:** Instituto de Biología Integrativa de Sistemas (I2SysBio), CSIC-Universitat de València, Valencia, 46980, Spain; Santa Fe Institute, Santa Fe, New Mexico, USA

**Keywords:** argonautes, host-virus interaction, mutualism, stress response, virus ecology and evolution

## Abstract

The heat shock response plays a role in the immune defense against viruses across various organisms. Studies on model organisms show that inducing this response prior to viral exposure enhances host resistance to infections, while deficient responses make individuals more vulnerable. Moreover, viruses rely on components of the heat shock response for their own stability and viral infections improve thermal tolerance in plants, giving infected individuals an advantage in extreme conditions, which aids the virus in replication and transmission. Here, we examine the interaction between the nematode *Caenorhabditis elegans* and its natural pathogen the Orsay virus (OrV) under heat stress. We found that OrV infection leads to differential gene expression related to heat stress, and infected populations showed increased resistance to heat shock. This resistance was associated with increased expression of argonautes *alg-1* and *alg-2*, which are crucial for survival after heat shock and for OrV replication. Overall, our study suggests an environmental-dependent mutualistic relationship between the worm and OrV, potentially expanding the worm’s ecological niche and providing the virus with extra opportunities for replication and adaptation to extreme conditions.

**IMPORTANCE:** Viruses have traditionally been viewed as self-serving pathogens that harm their hosts to ensure their own survival. However, recent studies are painting a different picture, revealing that viruses are ubiquitous and not always linked to diseases. In the realm of plant pathology, it has been long noted that the outcome of an infection hinges on environmental factors. Here, we show that the interaction between an animal virus and its natural host can be mutualistic under harsh temperature conditions. Our research highlights that this shift towards mutualism hinges on the expression of argonaute proteins.

---

The outcome of host-pathogen interactions strongly depends on environmental factors, with instances in which virulence is increased under abiotic stresses and situations in which more severe stresses result in reductions of pathogen’s prevalence (1). In the context of global warming, an environmental factor of particular relevant to better forecast the evolution and emergence of infectious diseases is thermal stress. Thermal stresses activate the heat shock response, a cellular mechanism that prevents proteotoxicity, and has diverse interactions with viral infections. Firstly, the heat shock response forms part of the antiviral immune response in a multitude of organisms, from plants to insects and mammals (2,3,4). Induction of a heat shock response in the nematode *Caenorhabditis elegans* and in mice prior to exposure to a virus increases the host resistance to the viral pathogen (5,6), whilst heat shock transcription factor deficient adult flies are hypersensitive to viral infection (7). Secondly, viruses require components of the heat shock response, such as Hsp90, for successfully completing their infectious cycle (8,9). Thirdly, viral infection has been reported to enhance thermal tolerance in some plants, conferring infected individuals an advantage over extreme conditions and enabling viruses to continue replicating and transmitting within the population (10).

In this study we examined the relationship between the nematode *C. elegans* and its natural parasite, the Orsay virus (OrV) (11), upon heat stress. We first looked into the transcriptional response elicited in infected worms, focusing on genes related to heat stress. For this purpose, we screened the differentially expressed genes upon OrV infection (12) for genes in the following GO-term categories: (*i*) cellular response to unfolded proteins, (*ii*) heat-shock binding proteins and (*iii*) response to heat. The three categories were overrepresented (Fisher’s exact test) among the significant differentially expressed genes (DEGs) upon OrV infection (Fig. 1A): 26.34% (*P* < 0.0001), 26.47% (*P* = 0.0003) and 13.16% (*P* = 0.0116), respectively. Among these DEGs, 60.71% were upregulated, suggesting that the transcriptional response induced by OrV has a strong overlap with the response elicited by a heat-shock. We then examined the ability of infected populations to withstand a semi-lethal heat-shock. As illustrated in Fig. 1B, we inoculated synchronized L1 populations of wild-type and JU1580 worms (the natural isolate in which OrV was first identified) with 4.92×10^8^ particles of OrV, a concentration that leads to activation of the *pals-5p::GFP* reporter (13) in ∼50% of the worms. Twenty-four hours post infection (hpi) we selected GFP positive worms, and 48 hpi we shifted the infected population from 20 to 37 ºC for 2 h. After the heat shock, we returned the worms to 20 ºC and 24 h later, we assessed the mortality for each treatment. Infected populations of both JU1580 and wild-type worms had a significant decrease in mortality upon heat-shock (Fig. 1C; Cohen’s *d* = −2.9574 □1.0070, *P* = 0.0033 and *d* = −4.0919 □1.8806, *P* = 0.0296, respectively), suggesting that OrV infection is able to confer thermal tolerance to *C. elegans*. Overall, averaging across worm genotypes, the observed infection-associated reduction in mortality due to heat-shock was largely significant (*d* = −3.5829 □1.1884, *P* = 0.0026).

**FIG 1.**
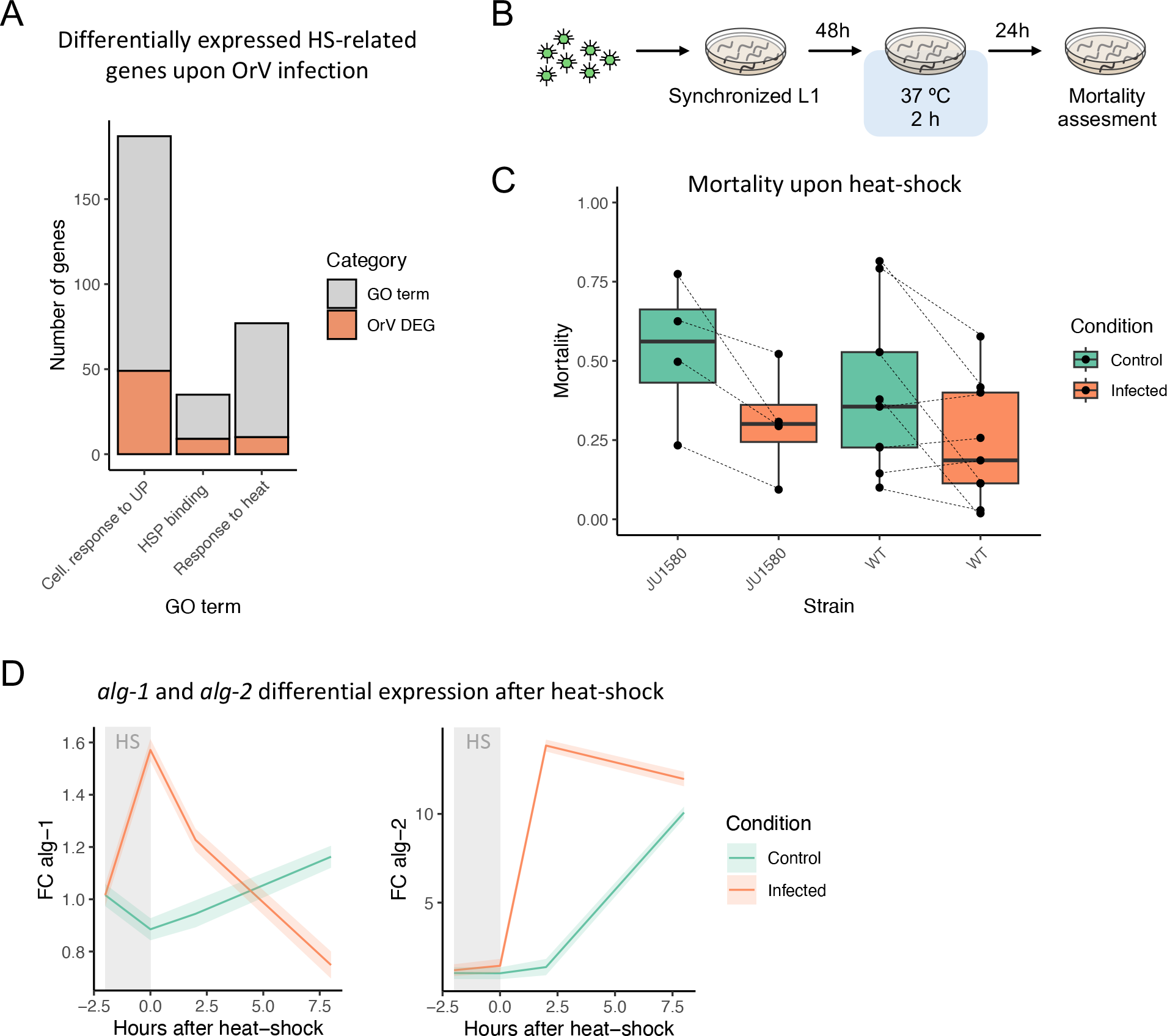
Interplay between OrV infection and heat-shock responses in *C. elegans*. (**A**) Significant enrichment of genes related to heat-shock responses that are associated to viral infection. (**B**) Schematic representation of the experiments performed. Four independent blocks were performed with the JU1518 strain and eight with the wild-type strain. (**C**) Effect of OrV infection in the mortality induced heat-shock. *P* values were obtained from the meta-analysis of experimental blocks as described in the Methods section. Gray lines connect mortality values of the same experimental block. (**D**) Analysis of the induction of two argonaute genes, *alg-1* and *alg-2*, upon heat-shock in control and OrV-infected worms. FC: fold-change in gene expression measured by RT-qPCR. Error bands represent □1 standard errors of the estimated marginal means.

Finally, in order to identify some of the players involved in this response we looked at the expression of the argonautes *alg-1* and *alg-2*, both of which are required for survival after heat-shock due to their role in microRNA biogenesis (14). Moreover, *alg-1* is necessary for OrV replication (15). Upon heat-shock, the expression of both of these genes increased significantly in infected versus control populations, showing quite different temporal dynamics (Fig 1D). In the case of *alg-1* (χ^2^ = 81.8560, 3 d.f., *P* < 0.0001), infection caused a sharp increase in its expression levels right after heat-shock, which decreased at 8 hours post heat-shock (hphs), at which point control worms displayed significantly higher expression levels of *alg-1*. In turn, infection caused a sharp increase in the expression levels of *alg-2* at 2 hphs and its levels remained high until 8 hphs (χ^2^ = 117.5381, 3 d.f., *P* < 0.0001), at which point the expression of *alg-2* also increased in control worms. These short but sharp increases in *alg-1* and *alg-2* levels may be critical in the increased thermal tolerance caused by OrV infection, although more studies would be necessary in order to fully elucidate the roles of these proteins in thermal tolerance and its interplay with viral infections.

Altogether, we showed here that infection of *C. elegans* by its natural virus OrV causes differential gene expression of many of the genes related to heat stress and confers protection against a mildly lethal heat-shock, both in wild-type worms and in the natural isolate JU1580. This protection may be at least partially explained by the upregulation of the argonautes *alg-1* and *alg-2*, which are required for survival after heat-shock and OrV replication. This mutualistic relationship between host and virus may have broad ecological consequences, expanding the host’s niche, allowing the virus to continue replicating and transmitting and thus having higher chances to adapt to even more extreme scenarios.

## METHODS

### Strain maintenance

Nematodes were maintained at 20 ºC on Nematode Growth Medium (NGM) plates seeded with *Escherichia coli* OP50 bacteria under standard conditions (16,17). ERT54 (*jyIs8 [pals-5p::GFP + myo-2p::mCherry]X* in an N2 Bristol background (13)) served as wild-type. JU2624, *(mjIs228[myo-2p::mcherry::unc54; lys-3p::eGFP::tbb-2]* in a JU1580 background) served as the natural isolate in which OrV was first identified. Both strains were obtained from Prof. M.A. Félix (11).

### OrV stock preparation and quantification

JU1580 worms were inoculated with OrV obtained from Prof. M.A. Félix (11), allowed to grow for 5 days and then resuspended in M9 buffer (0.22 M KH_2_PO_4_, 0.42 M Na_2_HPO_4_, 0.85 M NaCl, 0.001 M MgSO_4_), let stand for 15 min at room temperature, vortexed, and centrifuged for 2 min at 400 g. The supernatant was centrifuged twice at 21,000 g for 5 min and then passed through a 0.2 μm filter. RNA of the resulting viral stock was extracted using the Viral RNA Isolation kit (NYZ tech). The concentration of viral RNA was determined by RT-qPCR and normalized using a standard curve.

For the standard curve cDNA of OrV was obtained using Accuscript High Fidelity Reverse Transcriptase (Agilent) and reverse primers at the 3’ end of the virus (see Table 1 for primers). Approximately 1000 bp of the 3’ end of RNA1 and RNA2 were amplified using forward primers containing 20 bp coding the T7 promoter and DreamTaq DNA Polymerase (Thermo Fisher) (Table 1). The PCR products were gel purified using MSB Spin PCRapace (Invitek Molecular) and an *in vitro* transcription was performed using T7 Polymerase (Merck). The remaining DNA was then degraded using DNAse I (Life Technologies). RNA concentration was determined by NanoDrop (Thermo Fisher) and the number of molecules per μL was determined using the online tool EndMemo RNA Copy Number Calculator (https://endmemo.com/bio/dnacopynum.php).

**TABLE 1.**
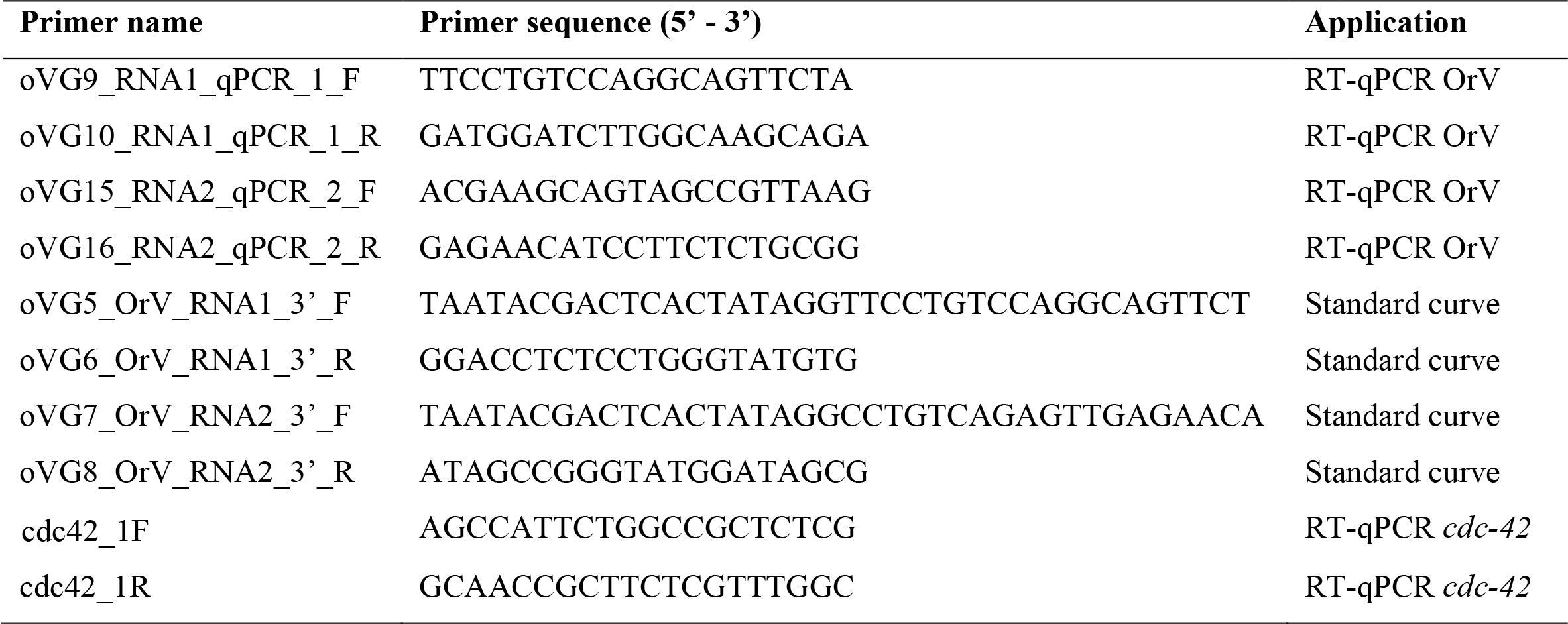
Sets of primers used for different purposes.

### Worm synchronization and OrV inoculation

In order to obtain synchronized worm populations, plates with eggs were carefully washed with M9 buffer in order to remove larvae and adults but leave the eggs behind. Plates were washed again using M9 buffer after 1 h to collect larvae hatched within that time span. Synchronized worm populations were inoculated with 4.92×10^8^ copies of OrV by pipetting the viral stock on top of the bacterial lawn containing the worms. Twenty-four hpi *pals-5p::GFP* negative worms were manually removed from the plates.

### Heat-shock

Synchronized population were shifted from a 20 ºC incubator to a 37 ºC water bath at 48 hpi for 2 h, after which they were returned to 20 ºC (Fig. 1B). Plates were sealed with parafilm and placed bottom down in order to ensure a quick temperature shift.

### Mortality assay

Twenty-four hphs mortality within the heat-shocked population was quantified. Worms were considered dead when they did not react to touch and had stopped pharyngeal pumping.

Mortality was evaluated in four and eight independent full blocks for JU1580 and wild-type strains, respectively. The number of plaques (biological replicates) within each experimental condition within blocks varied between 2 and 13 (median 3). Mean and standard deviation of mortality were estimated for each infection condition, worm strain and experimental block. Data were analyzed using a meta-analysis approach for continuous data, with infection status considered as a random factor, using Cohen’s *d* (□1 standard error) as a normalized estimation of the effect size, and restricted maximum-likelihood for parameter’s estimation. These analyses were done using SPSS version 29.0.1.1 (IBM Corp).

### Quantification of *alg-1* and *alg-2* expression

To quantify expression levels of *alg-1* and *alg-2*, wild-type control and infected worms were used. Worms were heat-shocked as described above and samples were taken before heat-shock, right after the heat-shock, and 2 and 8 hphs. Worms were collected with PBS 0.05% Tween and washed 3 times before freezing in liquid N_2_. RNA was extracted using Trizol as previously described (12). RT-qPCRs were performed using Power SYBR Green PCR Master Mix (Applied Biosystems) on an ABI StepOne Plus Real-time PCR System (Applied Biosystems). Ten ng of total RNA were loaded and samples were normalized to expression of *cdc-42*.

Fold-change values for each mutant genotype were independently fitted to a generalized linear model with a Normal distribution and identity function with infection status and hpsh incorporated as orthogonal random factors. The interaction between the two factors is reported in the text as an evaluation of the differences in temporal expression dynamics.

## ACKNOWLEDGEMENTS

We thank María J. Olmo-Uceda and Rubén González for helpful comments and discussions and Francisca de la Iglesia and Paula Agudo for excellent technical support.

This work is supported by grants PID2022-136912NB-I00 funded by MCIN/AEI/10.13039/501100011033 and by “ERDF a way of making Europe”, and CIPROM/2022/59 funded by Generalitat Valenciana to S.F.E. V.G.C. was supported by grant FJC2021-047264-I funded by MCIN/AEI/10.13039/501100011033 and by NextGenerationEU/PRTR.

